# Room-temperature fragment screening of soluble epoxide hydrolase by serial crystallography

**DOI:** 10.64898/2026.06.01.729266

**Authors:** Andreas Dunge, Gabrielle Wehlander, Gisela Brändén, Helena Käck

## Abstract

Room temperature serial crystallography offers advantages over conventional cryo-crystallography, such as simplified crystal handling and the possibility to avoid potential artefacts associated with cryo-trapping. However, to be considered as an alternative for drug discovery, where compound availability may be limited and speed of structure delivery is a key factor, it suffers from several limitations. To address these challenges, we have optimized a serial crystallography workflow for ligand soaking, data collection and data processing, significantly reducing time and reagent consumption to make it a viable option for drug discovery applications, herein exemplified by crystallographic fragment screening. Our approach incorporates the use of dried-in fragment cocktails on fixed target supports, compatible with 96-well plates for crystal soaking, and an efficient data processing pipeline tailored for serial crystallography. To validate our workflow, we conducted an in-crystal fragment screen at room temperature on the protein soluble epoxide hydrolase. The screen comprised 384 compounds and resulted in identification of 40 fragment binders corresponding to a hit rate of 10.4 %. The resulting room-temperature structures are of high quality and reveal opportunities for specific interaction within the highly hydrophobic active site of soluble epoxide hydrolase. Finally, we discuss potential avenues for further workflow optimization, highlighting the future potential of this approach for drug discovery.

**Synopsis:** We have developed a workflow that allowed us to efficiently conduct a fragment screen at room temperature using serial crystallography, of interest for future drug discovery campaigns.

## 1. Introduction

Over the last 20 years fragment-based drug discovery (FBDD) has developed into a well-established method for identification of high-quality lead compounds in drug discovery (Erlanson *et al*., 2016). A testimony of its success is the number of clinical candidates delivered based on chemical equity originating from fragment approaches (Woodhead *et al*., 2024). In addition, the recently approved AKT inhibitor, Capivasertib, developed for treatment of breast cancer is an example of a novel drug derived from a fragment-based project (Addie *et al*., 2013, Saxty *et al*., 2007, Caldwell *et al*., 2008).

The concept of FBDD is based on the idea that smaller compounds, so-called fragments (MW < 300 Da), can probe a binding pocket in a target protein for ideal interactions more efficiently compared to larger molecules that may be sterically restricted. Even though fragments are typically weak binders, it is possible to generate potent lead-like compounds that demonstrate optimal binding interactions with the target protein by merging and/or expanding fragment hits (Blundell *et al*., 2002, Whittaker *et al*., 2010, Blundell & Patel, 2004). Since the compound collections maintained by pharmaceutical companies largely are based on chemistry from prior projects, the evolution of fragments to lead compounds allows the creation of novel, typically under-represented chemistry. Thus, FBDD has been argued to be particularly important for unprecedented target classes (Fuller *et al*., 2016). In addition, the effective sampling of chemical space by fragments significantly reduces the number of compounds required for a screen. Typically fragment libraries contain in the range of 100-10000 compounds, compared to the millions of compounds that are tested in high-throughput screening settings (Kirsch *et al*., 2019, Bon *et al*., 2022).

Due to the weak binding of fragments, sometimes in the mM range, sensitive screening methods are required to identify the initial hits. Therefore, in addition to biophysical methods such as nuclear magnetic resonance (NMR) and surface plasmon resonance, protein crystallography is extensively used when screening for fragment hits (Blundell & Patel, 2004, Schiebel *et al*., 2016). Crystallography as a screening method for fragments offers several advantages. Firstly, soaking of protein crystals at high fragment concentrations has the potential to reveal very weak binders, not detectable by other methods (Schiebel *et al*., 2016, Davies *et al*., 2016). Secondly, the resulting structures reveal the binding pose of the hits, information which is often critical for fragment merging or expansion into more potent compounds.

For most fragment screens the structures are obtained through conventional X-ray crystallography where data is collected from a single crystal rotated in an X-ray beam while kept at cryogenic temperatures to avoid radiation damage (Hartshorn *et al*., 2005, Maveyraud & Mourey, 2020). This method is both well established and user friendly, but it does have limitations. It requires a robust protocol for producing large crystals, preferably suitable for soaking of compounds. Furthermore, the manual steps involved in soaking and harvesting crystals are labour intensive and limit the throughput. Therefore, multiple synchrotrons have developed platforms to facilitate the workflow, for example by including imaging protocols for crystal identification and subsequent use of acoustic dispensing of compounds. However, crystals still need to be harvested manually (Lima *et al*., 2020, Douangamath *et al*., 2021, Wollenhaupt *et al*., 2021).

The fact that the diffraction data from an X-ray crystallography fragment screen are typically collected at cryogenic temperature means that the resulting structural information is representative of the static frozen states. Interestingly, there are examples showing that cryo-trapping can lead to structural motifs being altered or hidden (Dunlop *et al*., 2005, Fischer *et al*., 2015). In addition, several studies have shown that room-temperature data show more defined density for flexible loop regions as well as alternate confirmations of side chains compared to the structures form cryo-cooled samples (Dods *et al*., 2017, Dunge *et al*., 2024, Uwangue *et al*., 2025). It has been suggested that the lack of density in the cryo-temperature structures is an artifact caused by the flash-cooling of crystals (Halle, 2004). Cryo-cooling of protein crystals was introduced as an efficient way to reduce secondary radiation damage by decreasing diffusion rates of deleterious radicals and other damaging species in the sample.

To overcome the limitations of cryo-crystallography, room-temperature diffraction data can be collected using a method called serial X-ray crystallography. In serial crystallography, thousands of randomly oriented micrometer-sized crystals (microcrystals) are presented to the X-ray beam and each crystal that interacts with the X-rays gives rise to one diffraction image, which can be merged into a complete dataset (Chapman *et al*., 2011). Serial synchrotron crystallography (SSX) is a method that is currently available at most synchrotrons (Weinert *et al*., 2017). Initially it was hampered by high sample consumption, due to the use of jets to flow the crystal sample across the X-ray beam (Weierstall, 2014). However, this has to a large extent been overcome by the use of so-called fixed-target devices, where a small amount of sample is loaded onto the sample support (Roedig *et al*., 2017). The support is subsequently translated across the X-ray beam so that each crystal is ideally exposed only once, resulting in a significant reduction of radiation damage compared to when room temperature diffraction data is collected by rotation of a single crystal. An added benefit of SSX is that the reliance on microcrystals makes it possible to use liquid handling tools, such as pipets and dispensing robotics when preparing crystal samples for data collection. Thereby the manual and labour intense step of crystal harvesting required for conventional X-ray crystallography where crystals are harvested, soaked and cryo-cooled may be avoided.

Soluble epoxide hydrolase (sEH) is a bifunctional enzyme catalyzing the conversion of epoxyeicosatrienoic acids (EETs) into dihydroxyeicosatrienoic acids (Fretland & Omiecinski, 2000). sEH consists of two domains that form a homodimer where the epoxide hydrolase activity resides in the C-terminal domain. In addition, phosphatase activity has been demonstrated for the N-terminal domain in-vitro, however the relevance of this activity is not clear (Newman *et al*., 2003). The active site catalyzing the hydrolase activity is large, mainly hydrophobic and composed of two distinct sub-pockets, referred to as the long branch or the left-hand side, and the short branch or the right-hand side, as shown in Figure 1 (Gomez *et al*., 2004). The pocket is constricted at the centre where the catalytic triad consisting of D335, H524 and D496 is located. Two tyrosine residues, Y466 and Y383 act as anchors and positions the substrate for catalysis. Inhibition of sEH leads to an accumulation of EETs which is postulated to have several positive effects on human health (Imig *et al*., 2002, Thomson *et al*., 2012) including reduction of hypertension, inflammation, and pain. As such, sEH constitutes an attractive target for drug discovery (Sun *et al*., 2021). Currently, there is one compound in Phase 1b trial for non-opiate pain reduction (Schmidt *et al*., 2024), however no compounds have been approved as drugs to date. A number of potent sEH inhibitors have been described, where the most common feature is a central urea or amide motif interacting with Y466, Y383 and D335 in the narrow central channel, flanked by hydrophobic motifs on either side (Figure 1). A recent review provides a comprehensive summary of sEH inhibitors (Sun *et al*., 2021). However, due to the hydrophobic nature of the binding pocket, it has proven challenging to develop inhibitors with properties suitable for clinical candidates including specific polar interactions. Fragment screening is well suited to identify hot spots in binding sites and has previously been applied to she (Xue *et al*., 2016, Amano *et al*., 2014), including one study that employed screening by X-ray crystallography (Amano *et al*., 2015). We therefore consider this target suitable for exploring the potential for room-temperature fragment screening using SSX.

**Figure 1.**
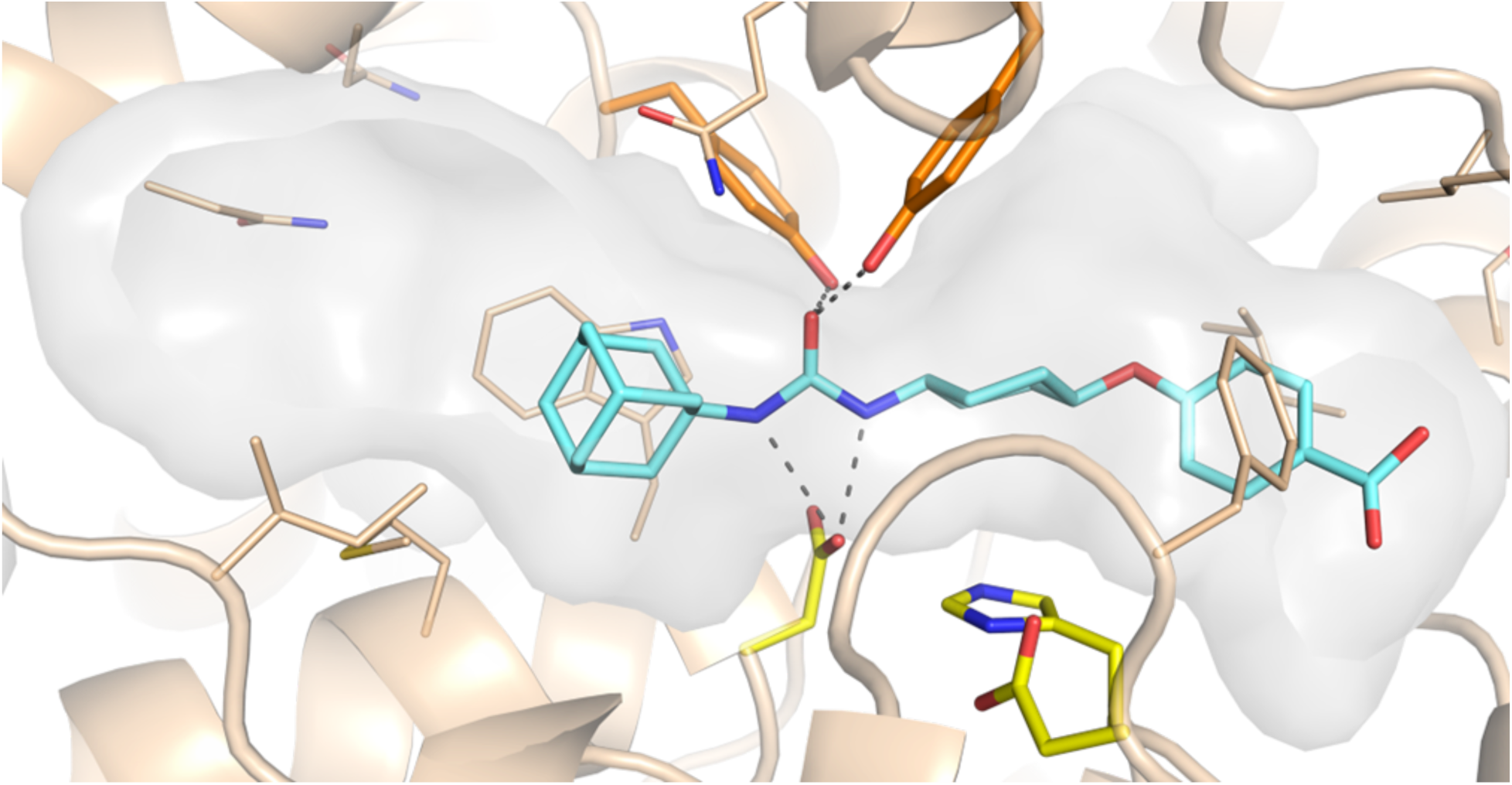
The structure of sEH in complex with the inhibitor *trans*-4-[4-(3-adamantan-1-yl-ureido)-cyclohexyloxy]-benzoic acid (pdb ID: 8QVH). The catalytic triad residues of are highlighted in yellow and the anchoring tyrosine residues in orange. The inhibitor is depicted in cyan.

In a previous study, we explored the use of SSX to determine structures of relevance for drug discovery (Dunge *et al*., 2024). Usings sEH as a model system, we developed a new workflow for screening and obtaining microcrystals which allowed us to determine the structure of six different compound complexes including fragments. All data were collected using a fixed-target serial crystallography setup at room temperature and resulted in structures of quality comparable with cryo crystallography. The study demonstrated the utility of SSX to obtain room-temperature fragment-bound structures at high resolution. In another recent study, a systematic comparison of fragment screening using room-temperature serial crystallography and single-crystal cryogenic data collection was conducted on the protein Fosfomycin-resistance protein A from Klebsiella pneumoniae (Gunther *et al*., 2025). Overall, more hits were found under cryogenic conditions, however the binding modes of the hits identified under both conditions were the same. The authors argued that hits found at both temperatures were less sensitive to thermal motion and disorder and therefore of higher relevance.

Herein, we describe the results from an X-ray crystallography fragment screen applied to sEH using room-temperature SSX. We present a workflow that combines the use of fixed-target supports with dried-in compounds for microcrystal soaking, allowing us to screen an X-ray library consisting of 384 diverse fragments, combined in 96 cocktails. Our study demonstrates that crystallographic fragment screening by SSX is highly feasible and that we obtain room-temperature structural information of excellent quality comparable with previous work using cryo-crystallography. Finally, future perspectives for the use of SSX for FBDD and drug discovery, including potential for further optimization, are discussed.

## 2. Methods

### 2.1. Protein expression, purification and crystallization

A C-terminally truncated sEH construct (residues 1-548) was expressed and purified as previously described (Xue *et al*., 2016). In brief, the protein was expressed in *Spodoptera frugiperda* (SF9) insect cells and purified using two interconnected HisTrap FF Crude (Cytvia) columns. Purified protein was stored in a buffer of 20 mM Tris-HCl pH 8, 150 mM NaCl, 1 mM TCEP and 10% glycerol, at a concentration of 13-20 mg/mL. sEH microcrystals were produced according to a previously established crystallization protocol (Dunge *et al*., 2024) by mixing protein (13-18 mg/mL) with a precipitant buffer (34 % (w/v) PEG3350, 0.1 M Tris-HCl (pH 8.5), 0.1 M Li_2_SO_4_) supplemented with 10 % v/v seed solution. in a 1:3 volume ratio. The crystals were produced by batch crystallization in 100 µL batches in Eppendorf tubes and prepared 12 to 48 hours prior to data collection.

### 2.2. Chemical Compounds

All fragments (1-10), listed in Table S1, are commercially available with CAS numbers: 198084-13-8, 41493-62-3, 146720-80-1, 28329-43-3, 87394-63-6, 292826-78-9, 1137-67-3, 57988-58-6, 28004-56-0, 128972-01-0.

### 2.3. Drying in fragment cocktails on fixed-target supports

The X-ray screen was performed using an in-house fragment library, tailor made for X-ray screening, of 384 compounds combined into 96 cocktails (Lucas *et al*., 2022). The cocktails were automatically dispensed on sheets holding 96 SerialFiX fixed-target supports with a 12.7 µm Kapton membrane (Ghosh *et al*., 2026) using a D300e Digital Dispenser (Tecan). The sheets of supports were placed on top of a Greiner 96-well microplate covered with parafilm to align the supports prior to dispensing the cocktails. For each sheet of 96 SerialFiX supports, eight different cocktails in six replicates were dispensed in rows. Every second row was left empty to allow folding of two supports together to sandwich the crystals between the membranes (Figure 2, step 1). 100 nL fragment cocktail at a concentration of 50 mM/fragment in DMSO was dispensed on the supports and left to dry overnight in a dust-free environment. The SerialFiX supports were prepared with dried compounds within a week of data collection.

**Figure 2.**
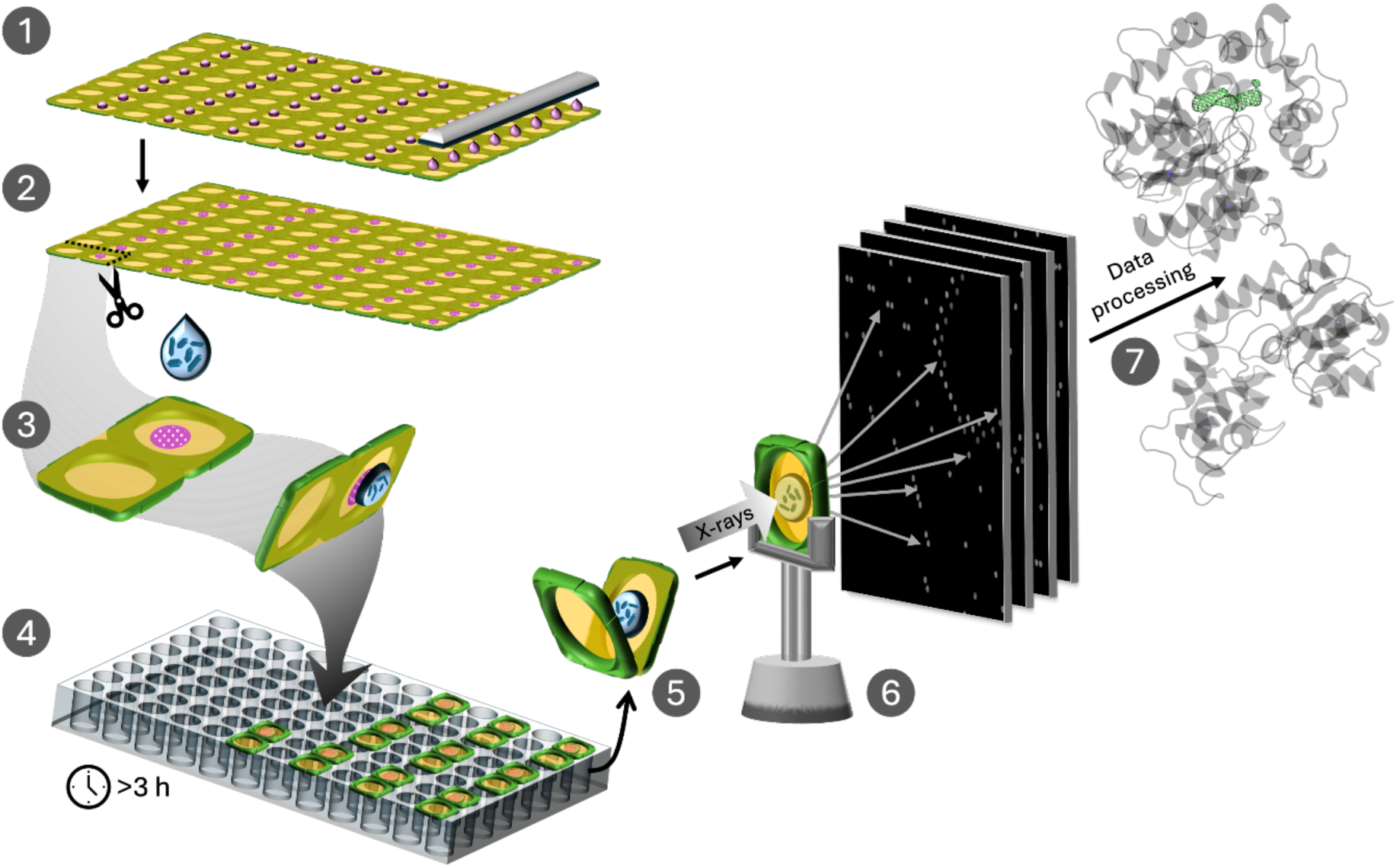
Optimized workflow for fragment screening using serial crystallography. (1) 100 nL fragment cocktail drops (pink) are automatically dispensed onto the membrane (yellow) of the SerialFiX fixed-target supports (green), with every second row left empty, and (2) allowed to dry. (3) To prepare for crystal soaking, arrays of 1×2 supports, with dried fragments on one support, are cut out, and a drop of concentrated crystals (blue) is added on top of the dried compound. (4) The crystals are soaked with the fragments by placing the supports on a 96-well plate containing a reservoir solution that mimics the mother liquor. (5) Prior to data collection, the 1×2 arrays of supports are folded onto each other and sealed using the adhesive tape (light green) to sandwich the crystals between the two membranes. (6) The enclosed supports are placed in a SerialFiX holder (grey) and mounted on the goniometer for data collection. (7) Data is processed to generate a model of the protein structure.

### 2.4. Soaking crystals on dried fragment cocktails

Each support with dried-in fragment cocktail was cut out together with its corresponding empty support, in 1×2 arrays. 1 µL concentrated microcrystals were thereafter added on top of the dried compounds, resulting in a concentration of 5 mM of each fragment in the sample. To prevent the crystal drops from dehydrating during soaking, the supports were placed on a Greiner 96-well microplate over a reservoir solution containing the precipitant buffer used for crystallisation, corrected for the protein buffer. The supports were sealed to the plates with the double-sided adhesive tape on the SerialFiX supports and crystals were soaked with the dried-in fragments during 3-10 hours prior to data collection. The process is schematically visualized in Figure 2.

### 2.5. Data collection and processing

SSX data were collected at the BioMAX beamline at MAX IV Laboratory using their fixed-target serial crystallography setup (Gonzalez *et al*., 2025). The 1×2 arrays of supports with fragment-soaked crystals were lifted from the soaking plate, folded on-top of each other to enclose the drop, attached to a sample-holder (serialx.se) and subsequently manually mounted on the goniometer in the beamline hutch. Data collection was monitored through the beamline control software *MXCuBE3* (Mueller *et al*., 2017), from where one or more mesh grids were defined and data collected through raster scans. All diffraction data were collected at room-temperature with an X-ray beam size of 20×20 µm with 100 % transmission, at a photon energy of 12.7 keV, a flux of 3-4×10^12^ photons s^−1^, and an exposure time of 11 ms on a Dectris Eiger 16M hybrid pixel detector.

All diffraction data were processed using programs within the *CrystFEL* suite, up to and including version 0.11.0 (White, 2019) at the MAX IV online cluster. Hit finding, indexing, and integration was conducted using *indexamajig*, with either of the peak-search algorithms *peakfinder8* or *peakfinder9* and the indexing algorithms *MOSFLM* (Powell, 1999) and *XDS* (Kabsch, 2010). The unit cell dimensions were determined manually with *cell_explorer*. Merging using scaling, partiality correction and post-refinement was conducted using *partialator* with the partiality model *xsphere*. The resolution limit for each dataset was determined based on signal-to-noise ratio >0.7, CC1/2 > 0.3, and completeness >95 %. To streamline the processing of multiple data set without much manual intervention, a script was used to manage the execution of the desired tasks.

### 2.6. Structure refinement

The structures were solved by the difference Fourier method using the structure 8QVM as the model (Dunge *et al*., 2024). Initially, a uniform refinement protocol was used for all structures starting with refinement using autoBUSTER version 2.11.8 (Bricogne *et al*., 2017), including an initial rigid-body refinement step. Subsequently the water molecules were updated such that existing waters with RMSD below 1σ in the 2FoFc map were removed and new ones were added based on having a signal stronger than 1σ in the 2FoFc map and 3σ in the FoFc map. Structures with resolution below 2.5 Å were excluded from further analysis, as these were considered of too low resolution for interpretation of ligand binding. A second round of refinement was performed to validate the newly added waters after which the fragments were fitted into the maps. Here the fragment in the cocktail best corresponding to the electron density were selected. In cases where the interpretation was ambiguous, all possible fragments were refined and the output analysed. Maps without any convincing difference density were classified as empty. After subsequent rounds of refinement, the structures were analysed such that a fragment was considered a hit if the 2FoFc had clear features corresponding to the fragment as well as interaction partners that supported binding. All model building was conducted in the software Coot (Emsley & Cowtan, 2004), using ligand constraints generated with WriteDict (Wlodek *et al*., 2006) or Grade (Smart *et al*., 2011) as necessary. Figures were prepared using PyMOL (*Schrodinger. The PyMOL Molecular Graphics System, Version 1.8.*, 2015).

## 3. Results and discussion

### 3.1. Choice of fixed target support

The Silson membrane (SiRN-5.0-200-2.5-1000) fixed-target supports used in our previous study (Dunge *et al*., 2024) had shown to be brittle and often broke during sample mounting. They were also prone to dehydration if left too long between crystal loading and data collection. Therefore, we explored the use of different supports. For this study the SerialFiX fixed-target devices with a 12.7 µm Kapton membrane (Ghosh *et al*., 2026) were selected based on their robustness and the fact that these are organized in a 12×8 array which is compatible with Greiner 96 well microplates. The latter facilitates compound soaking, as the supports can be mounted over plates containing reservoir solution to avoid dehydration of the crystals during the soaking process. It also simplifies visual inspection of the crystals in a microscope. Moreover, the compatibility with the 96-well plate format enables the use of automatic dispensing tools designed for plates.

### 3.2. Sample preparation

To optimize the workflow, we evaluated the possibility to dry-in fragments on the fixed-target membrane prior to adding crystals in order to facilitate soaking of crystals. This method has been described for compound introduction when working with large crystals in preparation for cryo-crystallography (Wollenhaupt *et al*., 2021) but has not been previously reported for fixed-target supports. Results from a pre-study confirmed that this method successfully allowed fragment binding.

Moreover, the compatibility of the SerialFiX devices with 96-well plates allowing addition of fragments using an automated dispensing system makes this an attractive approach. Thus, in advance of the beamtime, the SerialFiX supports were prepared with dried-in fragments from a fragment set dedicated to X-ray crystallography containing 384 fragments combined into 96 cocktails (Lucas *et al*., 2022). This was done by attaching the supports to a 96-well plate and subsequently dispensing 100 nL of each fragment cocktail using an automatic dispensing device (Figure 2, step 1-2). Every second row was left empty, so that it could be used as a lid when closing the support. The supports were then cut into 1×2 arrays where one of the membrane surfaces contained the dried-in fragment cocktail. We dispensed one cocktail per row, resulting in a total of 6 supports per cocktail, leaving a few spares in case some experiments needed to be repeated. The total consumed volume per cocktail including the dead volume in the dispenser was 2 µL.

At the beamline, crystal soaking with compounds was performed by adding 1 µL of concentrated crystal slurry, prepared as described previously (Dunge *et al*., 2024), to the membrane of each support that contained the dried-in fragments. The crystals were incubated for 3-10 h by placing the support over a reservoir mimicking the mother liquor (Figure 2, step 3-4). This efficiently prevents dehydration of the crystals during fragment soaking. In preparation for data collection, the supports were carefully removed from the plate, folded and sealed tightly using the adhesive tape that surrounds each membrane, and then placed in the SerialFiX holder for mounting on the goniometer (Figure 2, step 5-6). The procedure is simple and can easily be done on-the-fly at the beamline.

### 3.3. Data collection and processing

Room-temperature SSX data collection was carried out at the BioMAX beamline at the MAX IV Laboratory (Gonzalez *et al*., 2025) by a mesh scan using a beam size of 20×20 µm. The time from entering the hutch to mount the sample until the data collection was completed for one support varied between eight to twelve minutes, depending on the number of grids used (Figure S1). In our previous work, we established that 5000 indexable images resulted in a complete sEH dataset of good quality (Dunge *et al*., 2024). Here, we noted that collecting data from two supports with concentrated crystals would fulfill that requirement, leading to a data collection time of 20 minutes per structure, on average.

At the time of data collection, real-time data analysis was not available at the beamline, limiting the immediate assessment of data quality and number of diffraction images obtained. While the ambition was to collect data from two fixed-target supports per cocktail, this number was increased for some samples where it was observed that the crystal density was low or that the crystals had dried out during the soaking process. As a result, data were collected from two fixed-target supports per cocktail for 80 of the cocktails, whereas for the remaining 16, data was collected from three to four supports to ensure that a complete dataset was obtained. SSX data for the complete fragment screen were collected at three different occasions and the entire screen was performed in a total time of approximately 32 h.

All data processing was performed using the CrystFEL software (White, 2019). We developed an in-house pipeline for data processing allowing multiple datasets to be processed in a consecutive fashion, with minimal manual intervention (Figure S2). This allowed us to index the data for all 96 datasets within approximately 72 hours followed by merging the intensities within 48 hours. Despite the efficient pipeline, some procedures had to be conducted manually, such as determining the unit cell and resolution cut-off for each data set.

### 3.4. Resulting data sets

The X-ray crystallography fragment screen by SSX resulted in 96 datasets ranging from 1.8 – 4.3 Å resolution, with the average resolution being 2.23 Å (Table S2). This is similar to the reported resolution of sEH structures deposited in the protein data bank (the average resolution for 88 structures from the same space group is 2.24 Å). For each dataset, we collected approximately 30,000 images on average, of which 7100 were indexable, corresponding to a data collection hit-rate of 24 % (Table S2). Upon comparing the hit-rate between individual supports soaked with the same fragment cocktail, we note a large variation in the number of indexed images which likely depends on differences in crystal density between the samples. As expected, the resolution increases with the number of indexable images up to approximately 7000 images, after which it reaches a plateau (Figure 3). Our initial assumption that 5000 images would be required for a qualitative data set was thus reasonable, possibly slightly underestimated.

**Figure 3.**
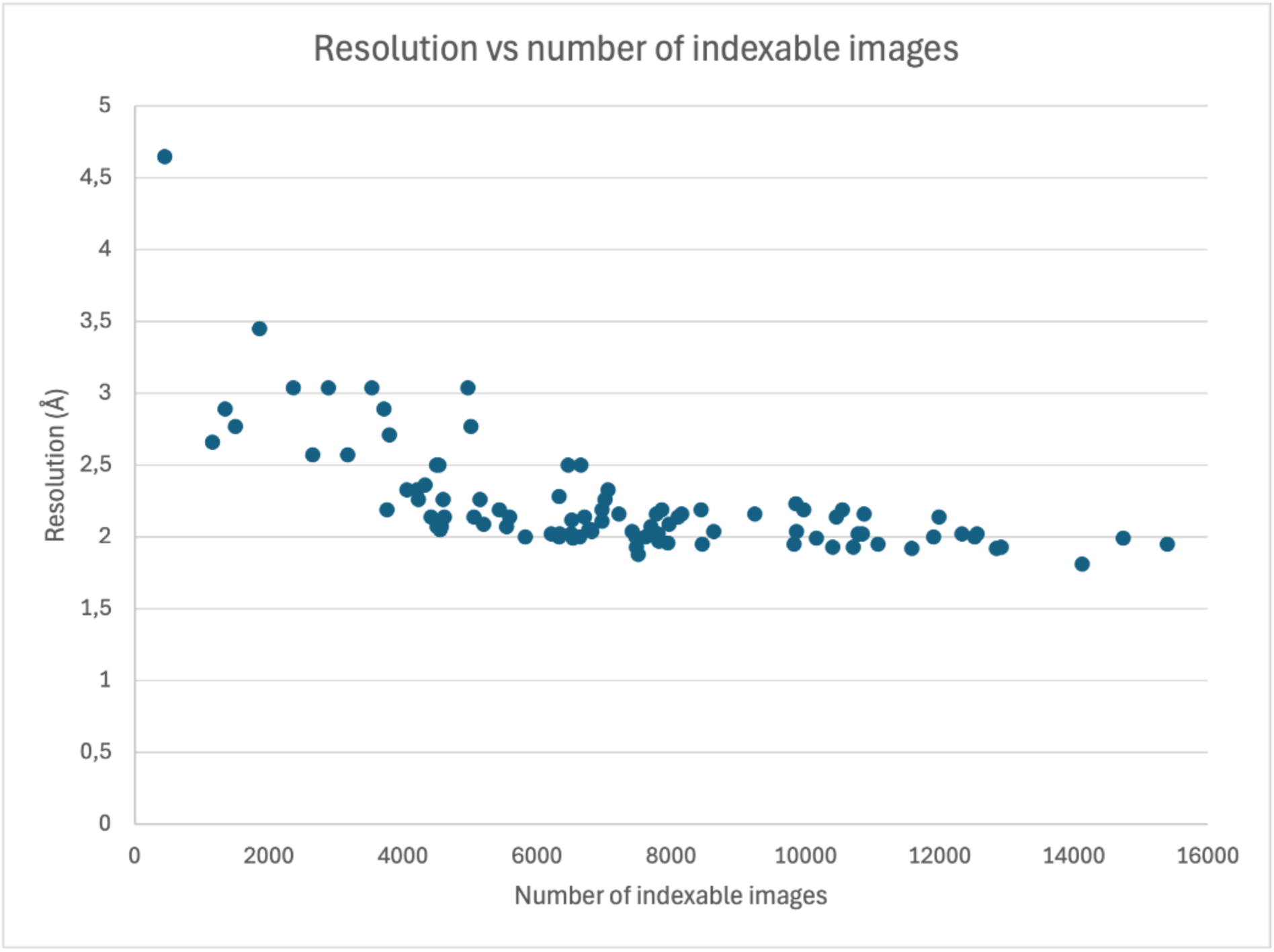
Number of indexable images per dataset plotted against the resolution. The resolution cut-off is based on a detector distance of 1.8 Å, CC1/2 > 0.3, signal-to-noise > 0.7 and completeness > 95%.

There is a clear trend that fewer indexable images results in lower resolution (Figure 3, Table S3). However, in one case a data set containing as little as 2650 indexed images results in a 2.6 Å resolution structure. The structures in the lowest resolution bin (2.51 - 4.3 Å) have a much-reduced hit rate (average of 13 %), suggesting that the diffraction quality is negatively affected in these samples. It is known that compound introduction can on occasions interfere with the crystal packing and thereby reduce the diffraction quality. Moreover, we have previously noted that the sEH crystals are very sensitive to dehydration. The issue of crystals drying out during data collection is largely avoided by using the SerialFiX chips. However, through visual inspection we noted a few cases where the sample showed signs of drying out, corresponding to data sets with a resolution lower than 2.5 Å.

Interestingly, fragments with low solubility did not seem to affect the data quality negatively. Figure S3 shows three droplets where the fragment cocktail did not fully dissolve in the crystallization buffer. Still, the number of indexable images and the resulting resolution is better than average for these three examples. The reason may be that when the support is closed the droplet is spread out so that the crystals cover the whole surface of the support whereas the undissolved fragments only cover a fraction of the total area. Therefore, our approach to dry-in the compounds prior to soaking appears to be an advantage in this context.

### 3.5. Overall outcome of the fragment screen

Structures were solved from all the resulting datasets, except for the two with lowest resolution (4.3 and 3.4 Å respectively). Those with a resolution lower than 2.5 Å were not considered suitable for interpretation of fragment binding and therefore not refined further, resulting in a total number of 82 structures being progressed. A fragment was considered a hit when the resulting density, after refinement, showed clear features corresponding to one of the fragments in the cocktail. Notably, several of the structures displayed electron density that could not be attributed to a specific fragment. This is not unusual when working with weak binders, that have been soaked at high concentrations. Partial occupancy in combination with overlapping binding poses can make it difficult to interpret the electron density and therefore distinguish individual fragments. In these cases, soaking the different fragments of the cocktail individually may have revealed further fragment binders. In summary, 33 of the refined structures contained 40 identifiable fragment hits, amounting to a total hit rate for the fragment screen of 10.4 %. Seven of the structures displayed binding of two independent fragments, for an example see Figure S4. In a previous crystallography-based fragment screen eight hits were reported from 800 screened fragments, displaying a significant lower hit-rate than we observed (Amano *et al*., 2015). Other cases of fragment screens have been described for sEH, however in these cases a high-concentration enzymatic assay in combination with NMR were used for initial screening, while crystallography was applied in the subsequent follow-up (Xue *et al*., 2016, Amano *et al*., 2014). Interestingly, for one of these cases an initial library of 4200 fragments resulted in 307 hits (7 %), however, only 41 % of the hits allowed successful structure determination, highlighting the benefit of using crystallography as a method for initial screening. The output from our room-temperature serial crystallography screen can also be compared to a recently published primary fragment screen targeting fosfomycin-resistance protein A where the fragment hit rate was 7.4 % using serial crystallography at room-temperature and as high as 32.6 % using single crystals at cryo temperature (Gunther *et al*., 2025).

A superposition of all structures classified as hits shows that the fragments efficiently probe the large substrate binding pocket in sEH (Figure 4a). Furthermore, it reveals that the lid of the binding pocket can adopt slightly different conformations (Figure 4b). This region displays a higher degree of flexibility in comparison with the rest of the protein and is most pronounced in the part of the pocket referred to as the short branch. This has previously been reported by others (Dotsch *et al*., 2024) and is consistent with the ability of sEH to bind a range of substrates. Interestingly, one of the two tyrosine residues that acts as anchors for the substrate, Y383, is part of the lid region. The flexibility of the lid allows this residue to adapt its position based on the size and position of the bound compound.

**Figure 4.**
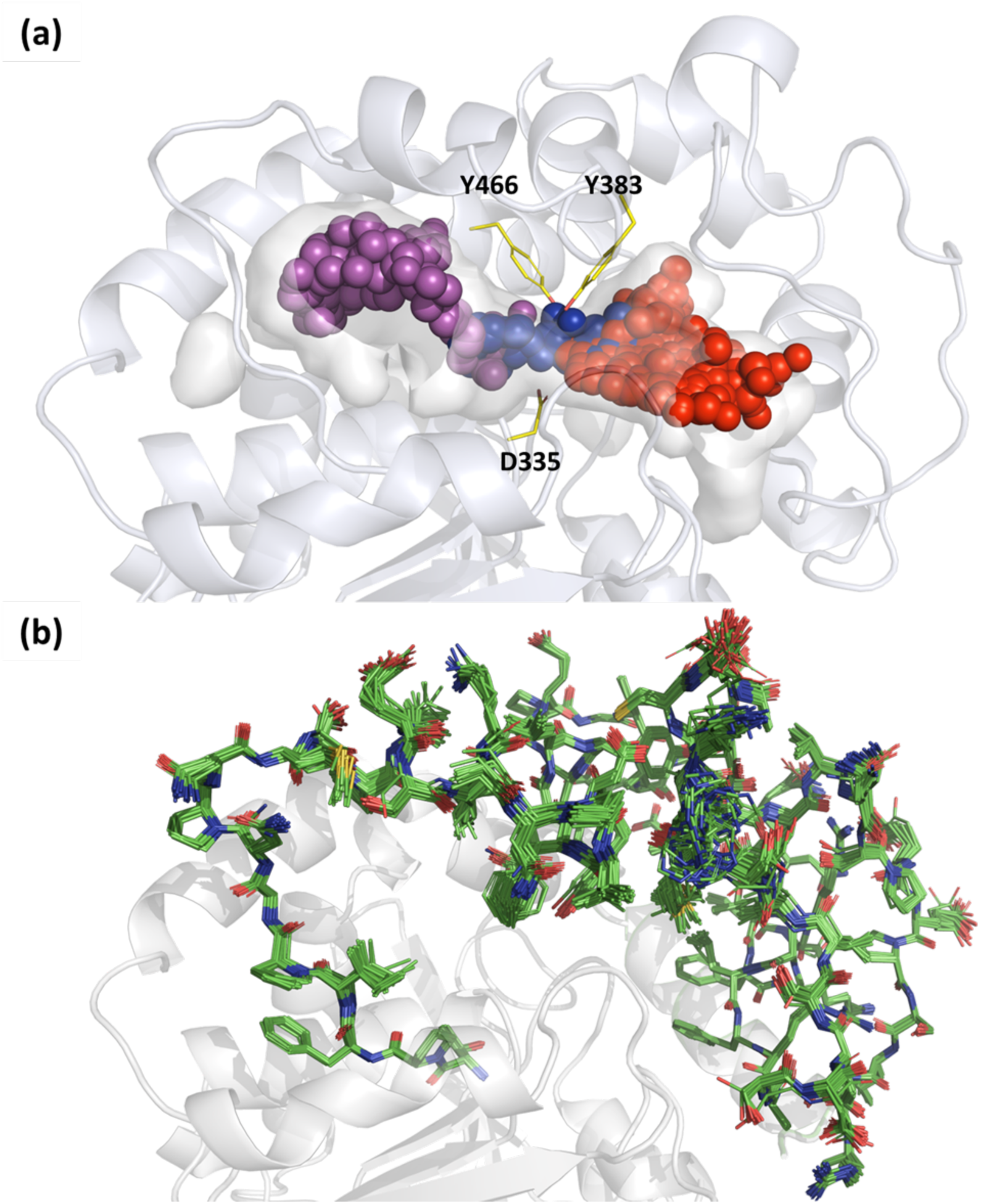
The catalytic site in the C-terminal domain of sEH. a) The colored spheres represent atoms from all fragments identified in the screen, superimposed. The tyrosine anchors (Y383, Y466) and the catalytic aspartate (D335) are depicted as sticks in yellow. Purple and red spheres represent fragments binding in the long respectively short branch of the pocket and blue spheres fragments binding to the constricted central part. b) A superposition of the lid region for all protein structures with confirmed binders. The fragments are excluded from the image for clarity.

The majority of the fragment hits, 28, are localized to the short branch of the sEH substrate pocket (red spheres, Figure 4a), while ten fragments bind to the long branch of the pocket (purple spheres, Figure 4a). Surprisingly, only two compounds were found to occupy the constricted central part of the pocket (blue spheres, Figure 4a). However, nine other fragments are within H-bonding distance to one or several of the substrate anchor residues Y383, Y466 or the catalytic residue D335 without spanning through the narrow central pocket. Interestingly, no fragments were found to bind outside the epoxidase catalytic pocket. A representative and diverse subset of the resulting fragments are discussed below (Figure 5).

**Figure 5.**
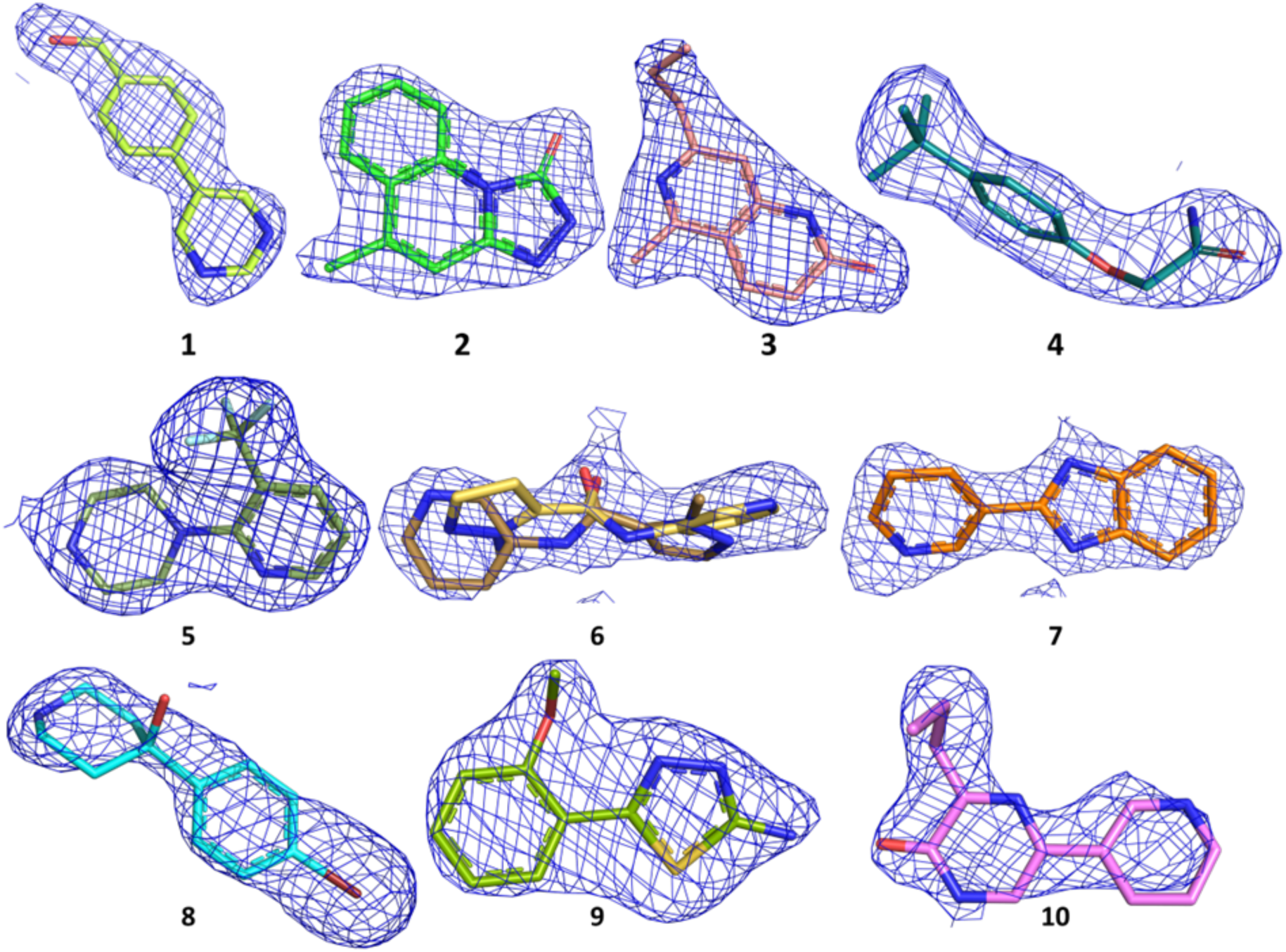
Selected fragment hits from the screen. The 2FoFc electron density is contoured at 1.0 σ (blue). Fragment **6** is modelled in two conformations.

### 3.6. Fragment hits in the short branch of the sEH binding pocket

The majority of the identified fragment-hits bind to the short branch of the sEH pocket. Similar to what has been observed previously, flat aromatic fragments with the ability to form a π-π interaction with the key residue H524 are favoured in this region (Eldrup *et al*., 2009), illustrated by fragments **1**-**3 (**Figure 6a-c). Interestingly, **4** stands out since it is positioned to form a potential edge to face interaction with H524 (Figure 6d). In contrast, **5** is an example of a fragment that does not form a π-π interaction with H524, instead its position is likely governed by the trifluoromethyl substituent which resides in a hydrophobic sub-pocket located near the lid region of the short branch defined by L428 and L408 (Figure 6e). A trifluoromethyl in this position has been described previously (Amano *et al*., 2014, Dotsch *et al*., 2024, Lee *et al*., 2014). Furthermore, **4**, **1** and **5** are examples of fragments forming specific interactions directly with the protein. Fragment **4** H-bonds with the main chain of V416, L417, S407 and L408, while **5** and **1** interact directly with the catalytic residues D496, Y383 and Y466. In addition, **1** forms a H-bond to the main chain of F497. A previous analysis (Bzowka *et al*., 2021) shows, not surprisingly, that many sEH binders interact with the catalytic residues, however interactions such as those made by **4** with V416 and S407 are rare. Several fragments, exemplified by **2** and **3,** lack direct specific interactions to the protein, however, a conserved water network provides opportunities for water mediated H-bonds to the protein (Figure 6b, c). Finally, we observe that some fragments bound in the short branch trigger the pocket to partially close by inducing a shift of F497 (Figure S5). This may potentially be due to a π-π interactions between the phenyl ring and the bound fragments. It has previously been proposed that F497 acts as a gate and regulates substrate access to the active site (Bzowka *et al*., 2021).

**Figure 6.**
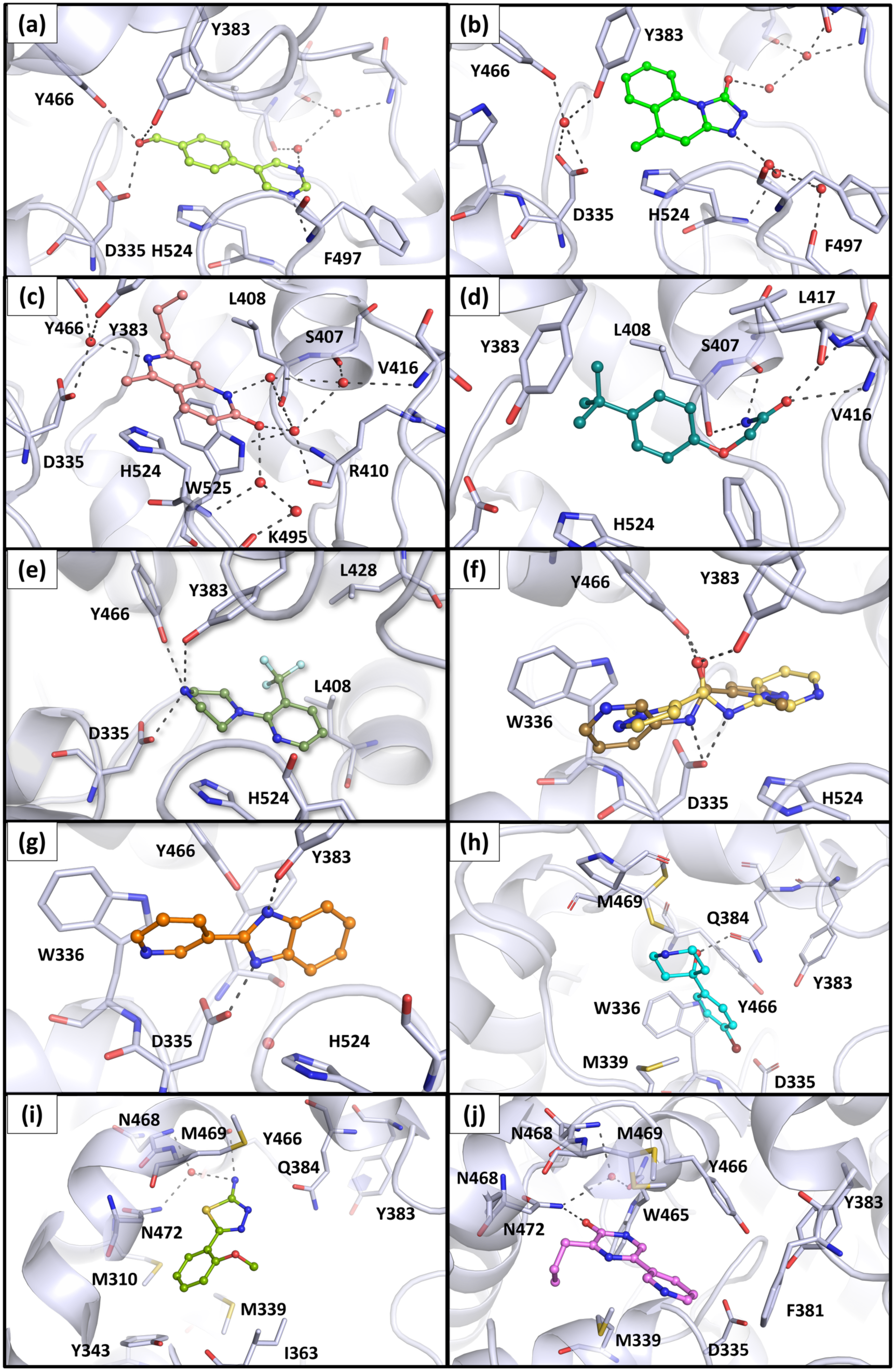
Representative fragment hits identified in the SSX screen bound to the active site of sEH. Panel a-j details interaction for **1-10** respectively and correspond to the following structures deposited in the protein data bank with the following IDs: 9QZA (**1**); 9QZR (**2**); 9QZS (**3**); 9QZU (**4**); 9QZV (**5**); 9QZW (**6, 10**); 9R02 (**7**); 9QZX (**8**); 9QZZ (**9**).

### 3.7. The central part of the binding pocket

Only two of the identified fragments hits, **6** and **7**, bind in the narrow, central part of the binding pocket which is characterized by the tyrosine anchors and the catalytic triad (Figure 6f, g). These two fragments interact directly with the key residues Y383, Y466 and D335, via an amide in the case of **6** and the benzimidazole moiety in the case of **7**. The former is common for many sEH inhibitors but also the benzimidazole motif has been described as binding to this site (Sun *et al*., 2021). Both fragments position an aromatic ring in the long branch, stacking against W336, while **6** also forms a π-π interaction with H524 in short branch. Nine additional fragments form H-bonds to at least one of the three central residues (Y383, Y466 or D335) without occupying the entire tunnel, e.g. **1** and **5 (**Figure 6a, e).

The low number of hits in the central part of the pocket is in stark contrast to the work published by Amano and coworkers, where 83 of 126 structurally characterized fragment hits bound to two out of three of Y383, Y466 or D335. Interestingly, the majority of those hits contained the already described urea and amide motifs. The fact that the primary screening was based on a biochemical assay, in combination with the use of a different fragment library, may account for this difference (Amano *et al*., 2014). Furthermore, in our previous SSX study we generated a set of ligand structures at room temperature and found that a PEG molecule was bound in this position in the apo structure (Dunge *et al*., 2024). This has not been observed previously and may be due to the increased PEG concentration used to grow microcrystals. Thus, competition with PEG may be the reason why we observe so few fragments with this binding mode. In fact, two fragment structures demonstrate simultaneous binding of a fragment and a PEG molecule (Figure S6).

### 3.8. Fragments hits in the long branch of the sEH binding pocket

The long branch of the pocket is large and characterized by hydrophobic residues, limiting the potential for specific interactions. Here we identified ten unique fragments, mainly located in two regions. The first site is located close to the active site as exemplified by **8 (**Figure 6h). Fragments binding at this site position an aromatic ring between the side chains of M339 and Q384, alternatively form a π-π interaction with W336. The Br atom of **8** is positioned close to the catalytic site within halogen bond distance to the catalytic residue D335. The second region is distant from the active site close to the amino acids M469, N472, I363 and Q384, exemplified by **9 (**Figure 6i). In addition to van der Waals interactions, this fragment forms a direct H-bond to the main chain of Y466 in addition to water mediated interactions to N468 and N472. Furthermore, we note that the side chain of M469 shifts to allow binding of the thiadiazole moiety of **9**. Finally, **10** is an example of a fragment anchored to the side chain of N472 similar to **9** but protruding into the pocket so that it overlaps with position of the piperidine moiety in **8** (Figure 6j).

In summary, the room-temperature fragment screen efficiently probed the sEH substrate binding pocket and resulted in a number of structurally divers hits. Among the identified hits are well-known motifs such as the amide interacting with the catalytic residues and the trifluoromethyl occupying a small hydrophobic pocket in the short branch. However, we also find binders such as **4** that display binding poses that are rarely observed for sEH inhibitors corroborating the value of fragment screening also for well-studied targets.

## 4. Conclusions and outlook

In this study, we have successfully optimized a serial crystallography workflow for ligand soaking, data collection and data processing. The focus was on reducing both the time required and reagent consumption for X-ray screening of compounds. As a proof-of-concept, we applied this workflow to conduct a room-temperature fragment screen on the target protein sEH. The resulting structures are of excellent quality with a resolution comparable to what has previously been reported for cryo-crystallography studies of the same protein. We observe a hit rate of 10.4 %, when considering only those fragments that can be clearly modelled into the electron density. Notably, the identified binders probe the binding pocket very efficiently. Consistent with other studies, our findings indicate that fragment screening successfully identifies potential for specific polar interactions, which may be valuable for modulating the property space of inhibitors binding to the predominantly hydrophobic ligand-binding pocket of she (Amano *et al*., 2014, Xue *et al*., 2016). A key distinction in our results is the large proportion of binders observed outside the narrow catalytic site, compared to a fragment screen based on an enzymatic assay (Amano *et al*., 2014), underscoring how different screening strategies can influence the hit distribution.

To device an effective workflow for ligand introduction, we leveraged the compatibility of fixed-target supports with the 96-well plate format. This enabled automatic dispensing of fragments to the supports and could potentially also be adopted for addition of crystal slurry onto the fixed-target supports, further streamlining the process. Moreover, the fact that the fragments were dried on the membranes prior to adding crystals, not only facilitates the workflow but also removes the potential issues associated with the presence of DMSO which can interfere with both crystal quality and ligand binding (Baker *et al*., 2020). In our previous study we were concerned that ligand precipitate would contribute to the background scattering and consequently reduce the signal-to-noise of the diffraction images. However, an unexpected benefit of drying-in the ligands on the membrane was that even if the ligand is not fully dissolved, it only affects a small section of the support. If needed, the affected area can easily be excluded during data collection. Each structure required 2 µL of protein crystal slurry and, to be compatible with automatic dispensing, 2 µL of 50 mM fragment solution. While this exceeds what is normally used for soaking a single crystal in fragment solution, it is comparable with the protein and ligand consumption required for co-crystallization of protein-ligand complexes and remains within an acceptable range. Importantly, all crystal handling was accomplished through pipetting eliminating the need for laborious, manual crystal manipulation.

Our approach demonstrates the feasibility of using SSX for drug discovery applications such as fragment screening where high throughput and short delivery times are crucial. However, to be a competitive method in terms of timelines, further optimization of the workflow is necessary. In this study the average time to collect a data set was 20 minutes, compared to 2-4 minutes for conventional cryo-crystallography, including mounting and automatic centering. The total time for a SSX dataset resulting from two supports encompasses manual mounting on the goniometer and definition of data collection parameters. One of the main limitations during our experiment was the need to manually exchange the sample, a task that is both labor intense and time consuming. Therefore, an obvious improvement would be to introduce sample changers compatible with the fixed-target supports. Such robotics are under development at several beamlines and have been made available at P09, DESY and PSI for a limited number of fixed-target supports. Ultimately, an automated data collection procedure without manual intervention, similar to what is available for cryo-crystallography, would be desirable. Even though SSX typically requires the collection of more images than conventional cryo-crystallography, state-of-the-art detectors with high frame rate combined with fast sample stages makes the actual data collection time less important in terms of reducing the experimental time (Zielinski *et al*., 2022). Finally, access to on-the-fly data analysis and direct feedback on hit rate is important to aid the assessment of data quality and obtain an initial structure within a reasonable time. An additional aspect not discussed in this paper is the issue of transporting room temperature crystals to the synchrotron for data collection (Ghosh *et al*., 2026). In our case we developed a crystallization protocol that allowed on-site crystallization which facilitated our experiments.

Our refined workflow showcases the viability of using room temperature serial crystallography as a tool for fragment screening and structure-based drug design, which together with on-going technical advances at synchrotrons have the potential to offer an attractive alternative to traditional cryo-crystallography techniques. Moreover, a simpler workflow will make the SSX technology more accessible for all types of studies, for example extensive exploration of the influence of temperature on structural features.

## Acknowledgements

X-ray diffraction data collection were carried out at the BioMAX beamline of MAX IV Laboratory (proposal number 20220186). We are grateful for assistance by the beamline staff and especially thank Monika Bjelcic at MAX IV Laboratory for help during data collection. The authors thank Anna Aagaard, Margareta Ek, Lisa Wissler and Linda Öster for fruitful discussions regarding ligand introduction.

## Author contributions

HK and GB conceived the experiment. Protein crystallization, X-ray diffraction data collection, X-ray diffraction data processing and refinement was performed by AD and GW. The manuscript was prepared through contributions of all authors. All authors have given approval to the final version of the manuscript.

## Data availability

The atomic coordinates and structure factors files for the crystal structures with compounds **1-10** bound have been deposited to the Protein Data Bank (www.pdb.org) with the following accession numbers: 9QZA (**1**), 9QZR (**2**), 9QZS (**3**), 9QZU (**4**), 9QZV (**5**), 9QZW (**6, 10**), 9R02 (**7**), 9QZX (**8**), 9QZZ (**9**).

## References

Addie, M., Ballard, P., Buttar, D., Crafter, C., Currie, G., Davies, B. R., Debreczeni, J., Dry, H., Dudley, P., Greenwood, R., Johnson, P. D., Kettle, J. G., Lane, C., Lamont, G., Leach, A., Luke, R. W., Morris, J., Ogilvie, D., Page, K., Pass, M., Pearson, S. & Ruston, L. (2013). J Med Chem 56, 2059–2073.

Amano, Y., Tanabe, E. & Yamaguchi, T. (2015). Bioorg Med Chem 23, 2310–2317.

Amano, Y., Yamaguchi, T. & Tanabe, E. (2014). Bioorg Med Chem 22, 2427–2434.

Baker, L. M., Aimon, A., Murray, J. B., Surgenor, A. E., Matassova, N., Roughley, S. D., Collins, P. M., Krojer, T., von Delft, F. & Hubbard, R. E. (2020). Commun Chem 3, 122.

Blundell, T. L., Jhoti, H. & Abell, C. (2002). Nat Rev Drug Discov 1, 45–54.

Blundell, T. L. & Patel, S. (2004). Curr Opin Pharmacol 4, 490–496.

Bon, M., Bilsland, A., Bower, J. & McAulay, K. (2022). Mol Oncol 16, 3761–3777.

Bricogne, G., Blanc, E., Brandl, M., Flensburg, C., Keller, P., Paciorek, W., Roversi, P., Sharff, A., Smart, O. S., Vonrhein, C. & Womack, T. O. (2017). Cambridge, United Kingdom: Global Phasing Ltd.

Bzowka, M., Mitusinska, K., Hopko, K. & Gora, A. (2021). Drug Discov Today 26, 1914–1921.

Caldwell, J. J., Davies, T. G., Donald, A., McHardy, T., Rowlands, M. G., Aherne, G. W., Hunter, L. K., Taylor, K., Ruddle, R., Raynaud, F. I., Verdonk, M., Workman, P., Garrett, M. D. & Collins, I. (2008). J Med Chem 51, 2147–2157.

Chapman, H. N., Fromme, P., Barty, A., White, T. A., Kirian, R. A., Aquila, A., Hunter, M. S., Schulz, J., DePonte, D. P., Weierstall, U., Doak, R. B., Maia, F. R., Martin, A. V., Schlichting, I., Lomb, L., Coppola, N., Shoeman, R. L., Epp, S. W., Hartmann, R., Rolles, D., Rudenko, A., Foucar, L., Kimmel, N., Weidenspointner, G., Holl, P., Liang, M., Barthelmess, M., Caleman, C., Boutet, S., Bogan, M. J., Krzywinski, J., Bostedt, C., Bajt, S., Gumprecht, L., Rudek, B., Erk, B., Schmidt, C., Homke, A., Reich, C., Pietschner, D., Struder, L., Hauser, G., Gorke, H., Ullrich, J., Herrmann, S., Schaller, G., Schopper, F., Soltau, H., Kuhnel, K. U., Messerschmidt, M., Bozek, J. D., Hau-Riege, S. P., Frank, M., Hampton, C. Y., Sierra, R. G., Starodub, D., Williams, G. J., Hajdu, J., Timneanu, N., Seibert, M. M., Andreasson, J., Rocker, A., Jonsson, O., Svenda, M., Stern, S., Nass, K., Andritschke, R., Schroter, C. D., Krasniqi, F., Bott, M., Schmidt, K. E., Wang, X., Grotjohann, I., Holton, J. M., Barends, T. R., Neutze, R., Marchesini, S., Fromme, R., Schorb, S., Rupp, D., Adolph, M., Gorkhover, T., Andersson, I., Hirsemann, H., Potdevin, G., Graafsma, H., Nilsson, B. & Spence, J. C. (2011). Nature 470, 73–77.

Davies, T. G., Wixted, W. E., Coyle, J. E., Griffiths-Jones, C., Hearn, K., McMenamin, R., Norton, D., Rich, S. J., Richardson, C., Saxty, G., Willems, H. M., Woolford, A. J., Cottom, J. E., Kou, J. P., Yonchuk, J. G., Feldser, H. G., Sanchez, Y., Foley, J. P., Bolognese, B. J., Logan, G., Podolin, P. L., Yan, H., Callahan, J. F., Heightman, T. D. & Kerns, J. K. (2016). J Med Chem 59, 3991–4006.

Dods, R., Bath, P., Arnlund, D., Beyerlein, K. R., Nelson, G., Liang, M., Harimoorthy, R., Berntsen, P., Malmerberg, E., Johansson, L., Andersson, R., Bosman, R., Carbajo, S., Claesson, E., Conrad, C. E., Dahl, P., Hammarin, G., Hunter, M. S., Li, C., Lisova, S., Milathianaki, D., Robinson, J., Safari, C., Sharma, A., Williams, G., Wickstrand, C., Yefanov, O., Davidsson, J., DePonte, D. P., Barty, A., Branden, G. & Neutze, R. (2017). Structure 25, 1461–1468.

Dotsch, L., Davies, C., Hennes, E., Schonfeld, J., Kumar, A., Guita, C., Ehrler, J. H. M., Hiesinger, K., Thavam, S., Janning, P., Sievers, S., Knapp, S., Proschak, E., Ziegler, S. & Waldmann, H. (2024). J Med Chem 67, 4691–4706.

Douangamath, A., Powell, A., Fearon, D., Collins, P. M., Talon, R., Krojer, T., Skyner, R., Brandao-Neto, J., Dunnett, L., Dias, A., Aimon, A., Pearce, N. M., Wild, C., Gorrie-Stone, T. & von Delft, F. (2021). J Vis Exp.

Dunge, A., Phan, C., Uwangue, O., Bjelcic, M., Gunnarsson, J., Wehlander, G., Kack, H. & Branden, G. (2024). IUCrJ 11, 831–842.

Dunlop, K. V., Irvin, R. T. & Hazes, B. (2005). Acta Crystallogr D Biol Crystallogr 61, 80–87.

Eldrup, A. B., Soleymanzadeh, F., Taylor, S. J., Muegge, I., Farrow, N. A., Joseph, D., McKellop, K., Man, C. C., Kukulka, A. & De Lombaert, S. (2009). J Med Chem 52, 5880–5895.

Emsley, P. & Cowtan, K. (2004). Acta Crystallogr D Biol Crystallogr 60, 2126–2132.

Erlanson, D. A., Fesik, S. W., Hubbard, R. E., Jahnke, W. & Jhoti, H. (2016). Nat Rev Drug Discov 15, 605–619.

Fischer, M., Shoichet, B. K. & Fraser, J. S. (2015). Chembiochem 16, 1560–1564.

Fretland, A. J. & Omiecinski, C. J. (2000). Chem Biol Interact 129, 41–59.

Fuller, N., Spadola, L., Cowen, S., Patel, J., Schonherr, H., Cao, Q., McKenzie, A., Edfeldt, F., Rabow, A. & Goodnow, R. (2016). Drug Discov Today 21, 1272–1283.

Ghosh, S., Banacore, A., Norder, P., Bjelcic, M., Kabbinale, A., Nileshwar, P., Wehlander, G., de Sanctis, D., Basu, S., Orleans, J., Vallejos, A., Chavas, L. M. G., Neutze, R. & Branden, G. (2026). J Appl Crystallogr 59.

Gomez, G. A., Morisseau, C., Hammock, B. D. & Christianson, D. W. (2004). Biochemistry 43, 4716–4723.

Gonzalez, A., Krojer, T., Nan, J., Bjelcic, M., Aggarwal, S., Gorgisyan, I., Milas, M., Eguiraun, M., Casadei, C., Chenchiliyan, M., Jurgilaitis, A., Kroon, D., Ahn, B., Ekstrom, J. C., Aurelius, O., Lang, D., Ursby, T. & Thunnissen, M. (2025). J Synchrotron Radiat 32, 779–791.

Gunther, S., Fischer, P., Galchenkova, M., Falke, S., Reinke, P. Y. A., Thekku Veedu, S., Rodrigues, A. C., Senst, J., Elinjikkal, D., Gumprecht, L., Meyer, J., Chapman, H. N., Barthelmess, M. & Meents, A. (2025). Nat Commun 16, 9089.

Halle, B. (2004). Proc Natl Acad Sci U S A 101, 4793–4798.

Hartshorn, M. J., Murray, C. W., Cleasby, A., Frederickson, M., Tickle, I. J. & Jhoti, H. (2005). J Med Chem 48, 403–413.

Imig, J. D., Zhao, X., Capdevila, J. H., Morisseau, C. & Hammock, B. D. (2002). Hypertension 39, 690–694.

Kabsch, W. (2010). Acta Crystallogr D Biol Crystallogr 66, 125–132.

Kirsch, P., Hartman, A. M., Hirsch, A. K. H. & Empting, M. (2019). Molecules 24.

Lee, K. S., Liu, J. Y., Wagner, K. M., Pakhomova, S., Dong, H., Morisseau, C., Fu, S. H., Yang, J., Wang, P., Ulu, A., Mate, C. A., Nguyen, L. V., Hwang, S. H., Edin, M. L., Mara, A. A., Wulff, H., Newcomer, M. E., Zeldin, D. C. & Hammock, B. D. (2014). J Med Chem 57, 7016–7030.

Lima, G. M. A., Talibov, V. O., Jagudin, E., Sele, C., Nyblom, M., Knecht, W., Logan, D. T., Sjogren, T. & Mueller, U. (2020). Acta Crystallogr D Struct Biol 76, 771–777.

Lucas, S. C. C., Borjesson, U., Bostock, M. J., Cuff, J., Edfeldt, F., Embrey, K. J., Eriksson, P. O., Gohlke, A., Gunnarson, A., Lainchbury, M., Milbradt, A. G., Moore, R., Rawlins, P. B., Sinclair, I., Stubbs, C. & Storer, R. I. (2022). RSC Med Chem 13, 1052–1057.

Maveyraud, L. & Mourey, L. (2020). Molecules 25.

Mueller, U., Thunnissen, M., Nan, J., Eguiraun, M., Bolmsten, F., Milàn-Otero, A., Guijarro, M., Oscarsson, M., de Sanctis, D. & Leonard, G. A. (2017). Synchrotron Radiation News 30, 22–27.

Newman, J. W., Morisseau, C., Harris, T. R. & Hammock, B. D. (2003). Proc Natl Acad Sci U S A 100, 1558–1563.

Powell, H. R. (1999). Acta Crystallogr D Biol Crystallogr 55, 1690–1695.

Roedig, P., Ginn, H. M., Pakendorf, T., Sutton, G., Harlos, K., Walter, T. S., Meyer, J., Fischer, P., Duman, R., Vartiainen, I., Reime, B., Warmer, M., Brewster, A. S., Young, I. D., Michels-Clark, T., Sauter, N. K., Kotecha, A., Kelly, J., Rowlands, D. J., Sikorsky, M., Nelson, S., Damiani, D. S., Alonso-Mori, R., Ren, J., Fry, E. E., David, C., Stuart, D. I., Wagner, A. & Meents, A. (2017). Nat Methods 14, 805–810.

Saxty, G., Woodhead, S. J., Berdini, V., Davies, T. G., Verdonk, M. L., Wyatt, P. G., Boyle, R. G., Barford, D., Downham, R., Garrett, M. D. & Carr, R. A. (2007). J Med Chem 50, 2293–2296.

Schiebel, J., Radeva, N., Krimmer, S. G., Wang, X., Stieler, M., Ehrmann, F. R., Fu, K., Metz, A., Huschmann, F. U., Weiss, M. S., Mueller, U., Heine, A. & Klebe, G. (2016). ACS Chem Biol 11, 1693–1701.

Schmidt, W. K., Cortes-Puch, I., McReynolds, C. B., Croston, G. E., Hwang, S. H., Yang, J., Pedersen, T. L., Wagner, K. M., Pham, T. T., Hunt, T. & Hammock, B. D. (2024). Clin Transl Sci 17, e70033.

Schrodinger. The PyMOL Molecular Graphics System, Version 1.8. (2015).

Smart, O. S., Womack, T. O., Sharff, A., Flensburg, C., Keller, P., Paciorek, W., Vonrhein, C. & Bricogne, G. (2011). Cambridge: Global Phasing Ltd.

Sun, C. P., Zhang, X. Y., Morisseau, C., Hwang, S. H., Zhang, Z. J., Hammock, B. D. & Ma, X. C. (2021). J Med Chem 64, 184–215.

Thomson, S. J., Askari, A. & Bishop-Bailey, D. (2012). Int J Vasc Med 2012, 605101.

Uwangue, O., Glerup, J., Dunge, A., Bjelcic, M., Wehlander, G. & Branden, G. (2025). Arch Biochem Biophys 769, 110419.

Weierstall, U. (2014). Philos Trans R Soc Lond B Biol Sci 369, 20130337.

Weinert, T., Olieric, N., Cheng, R., Brunle, S., James, D., Ozerov, D., Gashi, D., Vera, L., Marsh, M., Jaeger, K., Dworkowski, F., Panepucci, E., Basu, S., Skopintsev, P., Dore, A. S., Geng, T., Cooke, R. M., Liang, M., Prota, A. E., Panneels, V., Nogly, P., Ermler, U., Schertler, G., Hennig, M., Steinmetz, M. O., Wang, M. & Standfuss, J. (2017). Nat Commun 8, 542.

White, T. A. (2019). Acta Crystallogr D Struct Biol 75, 219–233.

Whittaker, M., Law, R. J., Ichihara, O., Hesterkamp, T. & Hallett, D. (2010). Drug Discov Today Technol 7, e147–202.

Wlodek, S., Skillman, A. G. & Nicholls, A. (2006). Acta Crystallogr D Biol Crystallogr 62, 741–749.

Wollenhaupt, J., Barthel, T., Lima, G. M. A., Metz, A., Wallacher, D., Jagudin, E., Huschmann, F. U., Hauss, T., Feiler, C. G., Gerlach, M., Hellmig, M., Forster, R., Steffien, M., Heine, A., Klebe, G., Mueller, U. & Weiss, M. S. (2021). J Vis Exp.

Woodhead, A. J., Erlanson, D. A., de Esch, I. J. P., Holvey, R. S., Jahnke, W. & Pathuri, P. (2024). J Med Chem 67, 2287–2304.

Xue, Y., Olsson, T., Johansson, C. A., Oster, L., Beisel, H. G., Rohman, M., Karis, D. & Backstrom, S. (2016). ChemMedChem 11, 497–508.

Zielinski, K. A., Prester, A., Andaleeb, H., Bui, S., Yefanov, O., Catapano, L., Henkel, A., Wiedorn, M. O., Lorbeer, O., Crosas, E., Meyer, J., Mariani, V., Domaracky, M., White, T. A., Fleckenstein, H., Sarrou, I., Werner, N., Betzel, C., Rohde, H., Aepfelbacher, M., Chapman, H. N., Perbandt, M., Steiner, R. A. & Oberthuer, D. (2022). IUCrJ 9, 778–791.

