## Supplementary material for "Room-temperature fragment screening of soluble epoxide hydrolase by serial crystallography"

**This PDF file includes:**
**Supplementary Figures, S1-S6**

**Supplementary Tables S1-S3**

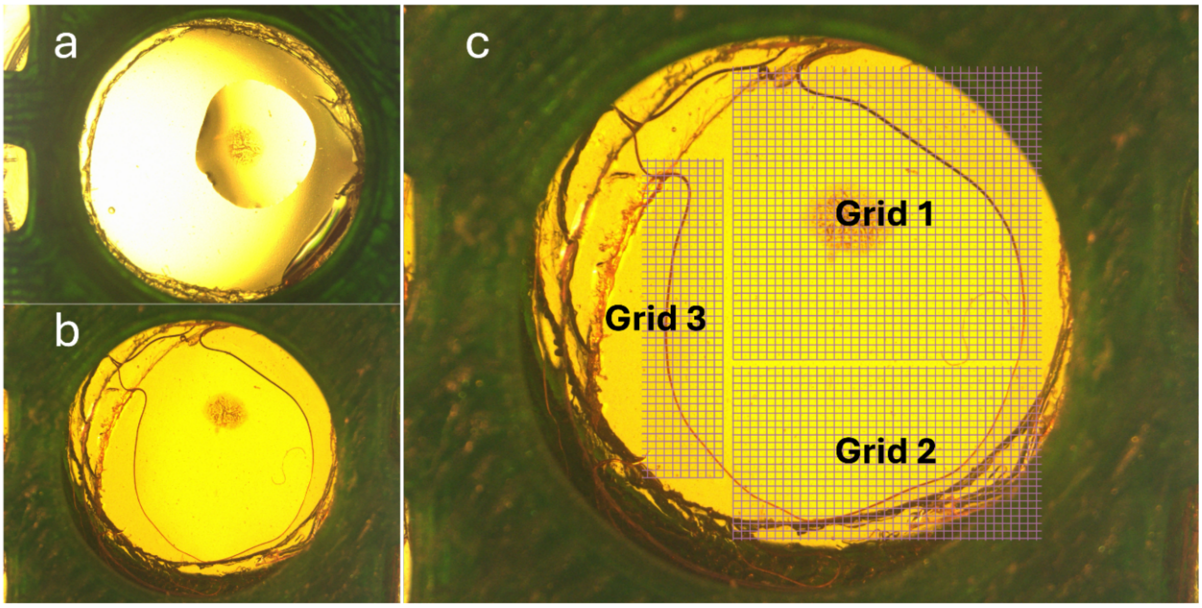

**Figure S1** Example of the sample distribution on the SerialFiX support, here demonstrated using only crystallization buffer (without crystals). A) 1 µL crystallization buffer is added on top of the dried-in fragments. B) The size and spread of the sample after closure of the fixed-target support. C) Example of how the grids for data collection could be defined.

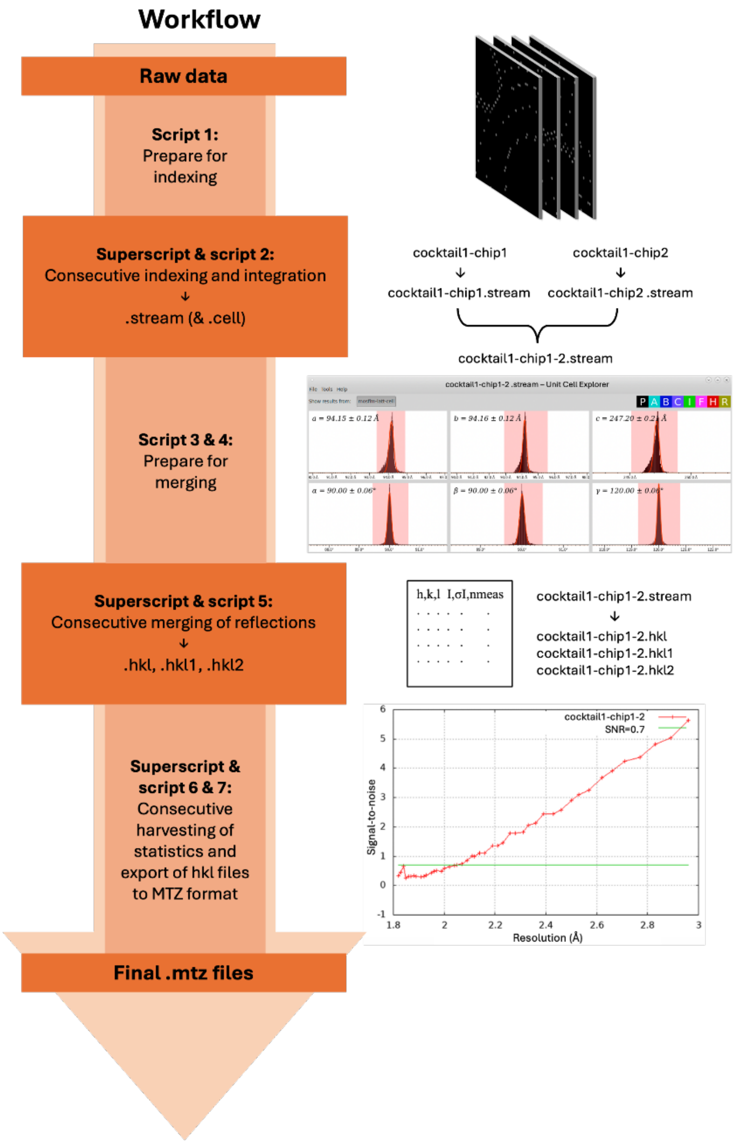

### What

**Script 1: Prepare for indexing**

i) Creates processing folders for all SerialFiX support collected

ii) Creates lists of the raw data files (diffraction images) to process

iii) Adds files and scripts needed for processing (geometry file, mask file, unit cell file, script 2)

**Script 2: Indexing and integration**

i) Indexes and integrates diffraction spots from the raw data files using CrystFEL's *indexamajig*.

ii) Integrated diffraction spots are stored in stream files (from which the unit cell dimensions can be manually determined using CrystFEL's *cell_explorer*)

**Script 3 & 4: Prepare for merging**

i) Merges streamfiles from the same cocktail (collected on separate SerialFiX supports)

ii) Creates new processing folders for the merged streamfiles and adds scripts necessary for further processing

**Script 5: Merging of reflections**

i) Merges reflections by scaling, partiality correction and post-refinement using CrystFEL's *partialator*

**Script 6: Statistics**

i) Compiles statistics from all datasets, using CrystFEL's *check_hkl* and *compare_hkl*

**Script 7: Export .hkl to .mtz**

i) Export reflection files ending with .hkl into MTZ format, using CrystFEL's *get_hkl*

**Superscript:** Automates the execution of a desired script located in subdirectories of the current directory. After finishing the execution of a desired script in one subdirectory, it progresses to initiate processing in the next subdirectory.

**Figure S2** Description of the software pipeline for data processing.

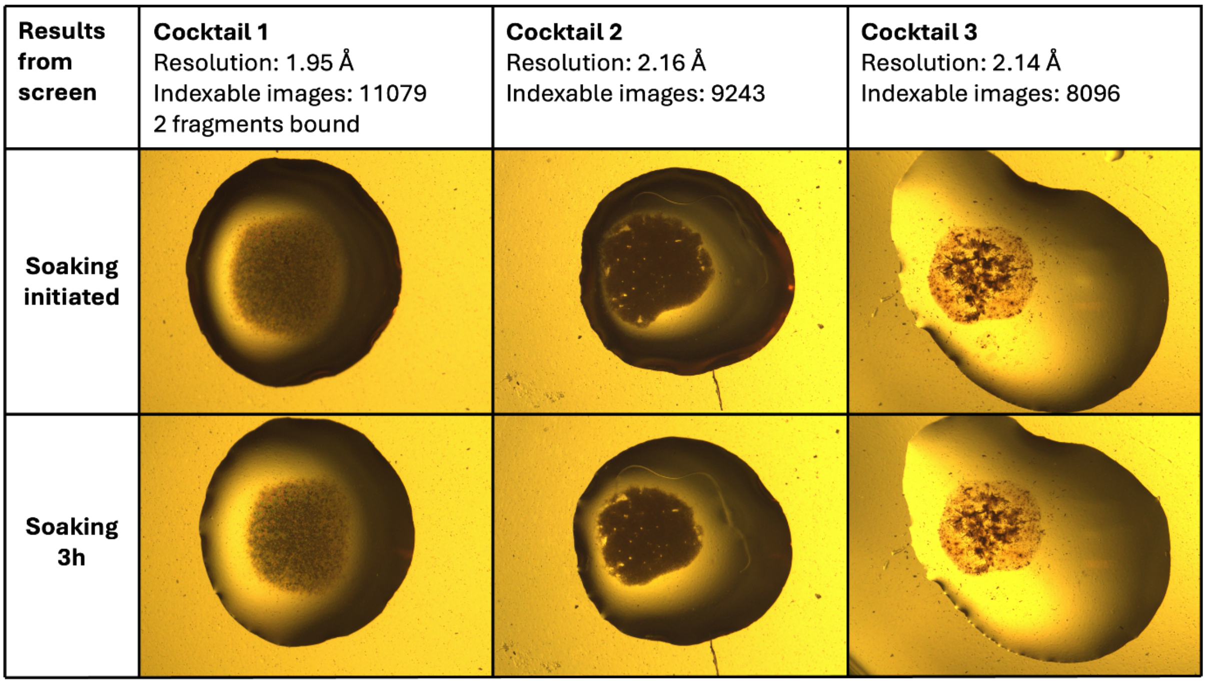

**Figure S3** Example of fragment cocktails that did not dissolve during soaking. The three cocktails resulted in data with a resolution better than the average resolution of 2.2 Å for all data sets. One of these resulted in a ligand-bound structure. For better visualization, the added drop contains only crystallization buffer and no crystals or protein.

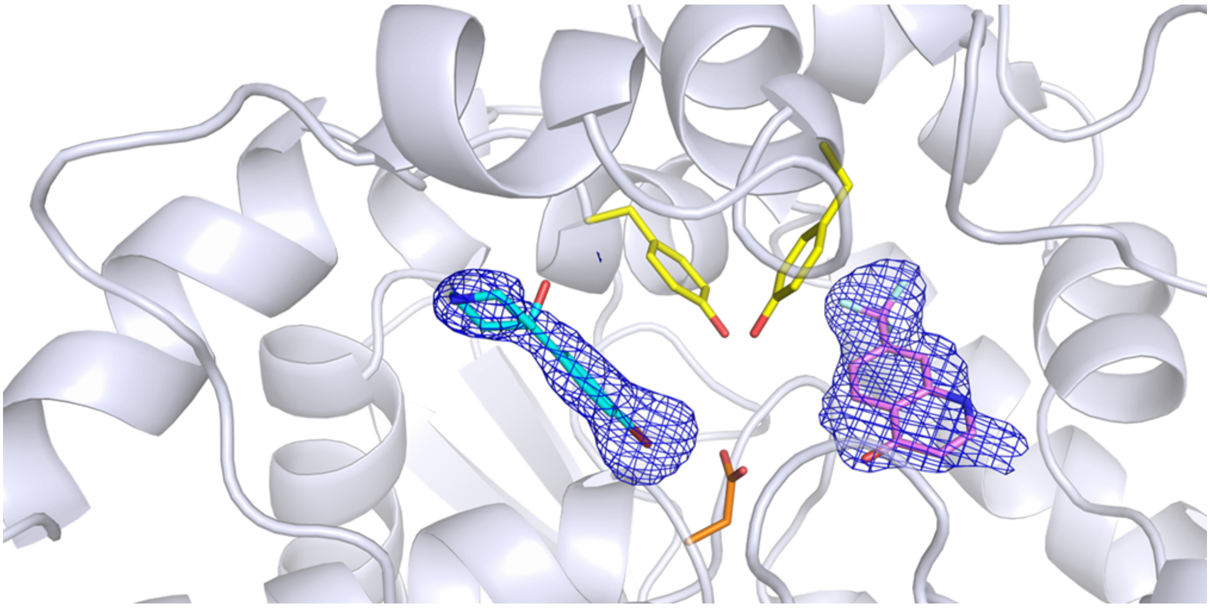

**Figure S4** An example of two fragments from the same cocktail binding simultaneously to two distinct sites in sEH. One fragment, **10**, is located in the large branch (cyan) and the other is in the short branch (pink, not discussed in the paper). The 2FoFc electron density map in blue is contoured at 1 σ.

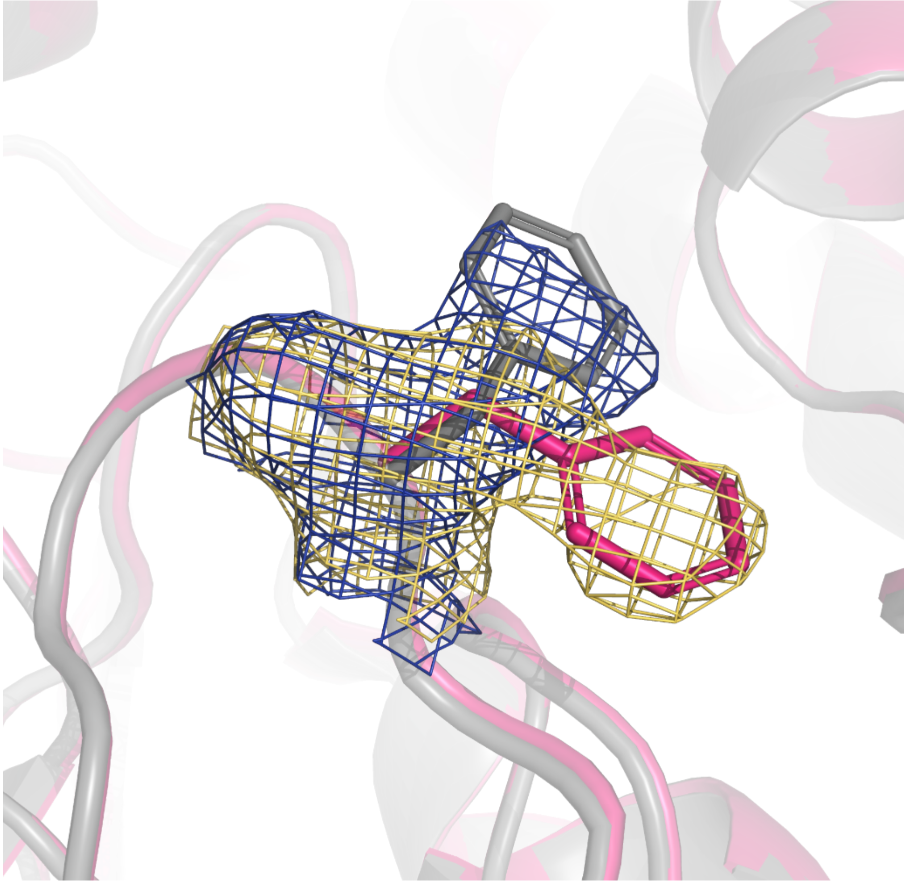

**Figure S5** Example of two structures with different conformation of the F497 side chain. The 2FoFc electron density map contoured at 1 σ displays F497 in the inwards conformation (map in blue and structure in grey) and the outward conformation (map in yellow and structure in pink), respectively.

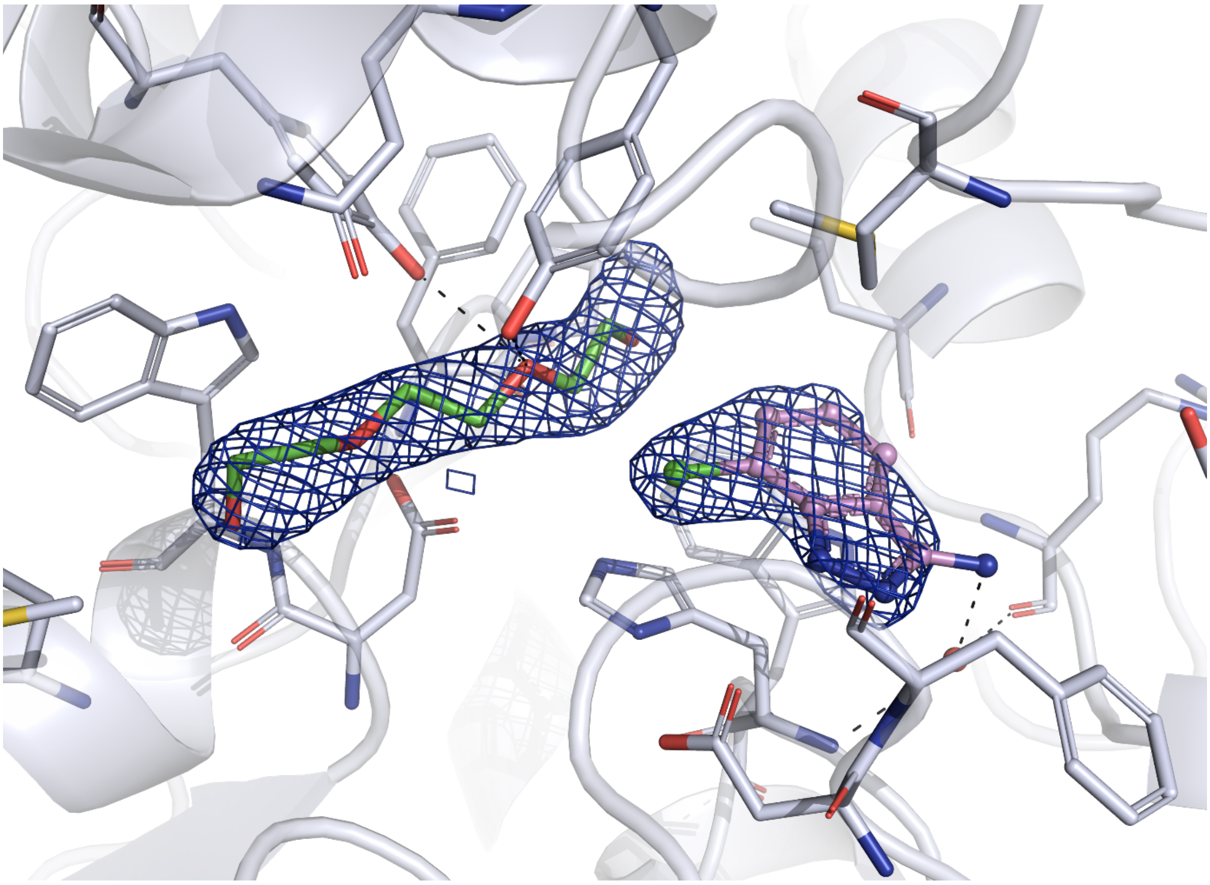

**Figure S6** Example of a PEG molecule binding to the central tunnel, while a fragment is bound in the short branch of the sEH active site. The 2FoFc electron density map (blue) is contoured at 1 σ.

**Table S1** Compounds discussed in the paper.

| Number in manuscript | SMILES | CAS number |
|---|---|---|
| 1 | c1cc(ccc1CO)c2cncnc2 | 198084-13-8 |
| 2 | Cc1cc2n[nH]c(=O)n2c3c1cccc3 | 41493-62-3 |
| 3 | CCCc1cc2c(ccc(=O)[nH]2)c(n1)C | 146720-80-1 |
| 4 | CC(C)(C)c1ccc(cc1)OCC(=O)N | 28329-43-3 |
| 5 | c1cc(c(nc1)N2CCNCC2)C(F)(F)F | 87394-63-6 |
| 6 | Cn1c(ccn1)C(=O)Nc2cccnc2 | 292826-78-9 |
| 7 | c1ccc2c(c1)[nH]c(n2)c3cccnc3 | 1137-67-3 |
| 8 | c1cc(ccc1C2(CCNCC2)O)Br | 57988-58-6 |
| 9 | COc1ccccc1c2nnc(s2)N | 28004-56-0 |
| 10 | CCCc1c(=O)[nH]cc(n1)c2cccnc2 | 128972-01-0 |

**Table S2** Data processing and refinement statistics. Values in parentheses are those for the highest-resolution shell.

| | Structure with fragment 1 | Structure with fragment 2 | Structure with fragment 3 |
|---|---|---|---|
| **Data collection** | | | |
| PDB id | 9QZA | 9QZR | 9QZS |
| Data collection date | Novemeber 2023 | April 2023 | April 2023 |
| Wavelength (Å) | 0.98 | 0.98 | 0.98 |
| Temperature (K) | 293 | 293 | 293 |
| a, b, c (Å) | 94.28, 94.28, 247.35 | 94.11, 94.11, 246.83 | 94.37, 94.37, 245.93 |
| α, β, γ (°) | 90, 90, 120 | 90, 90, 120 | 90, 90, 120 |
| Resolution range (Å) | 38.77-1.93 (1.93-1.96) | 123.42-1.85 (1.85-1.86) | 122.97-2.04 (2.04-2.05) |
| Total no. of reflections | 4782755 | 7027436 | 9016005 |
| No. of unique reflections | 92556 | 93486 | 69620 |
| Completeness (outer shell) (%) | 100 (100) | 100 (100) | 100 (100) |
| Redundancy (outer shell) | 51.67 (33.6) | 75.17 (50.7) | 129.50 (87.9) |
| I/σ(I) | 4.68 (0.88) | 6.16 (0.94) | 4.94 (1.00) |
| $CC_{1/2}$ | 0.97 (0.44) | 0.97 (0.64) | 0.96 (0.64) |
| $R_{split}$ (%) | 15.57 (114.60) | 13.30 (68.71) | 16.91 (69.17) |
| Wilson B factor ($Å^2$) | 34.24 | 37.13 | 47.75 |
| Total no. of images | 19273 | 35046 | 19175 |
| Indexed images | 7475 | 11584 | 6518 |
| No. of supports collected | 2 | 2 | 2 |
| Soaking time (h) | 5h | 5-5:30h | 6-6:30h |
| Fragment(s) modelled | 1 | 1 | 1 |
| **Structure refinement** | | | |
| Resolution range (Å) | 1.93-27.61 | 1.85 - 26.76 | 2.04-29.07 |
| No. of reflections used in refinement | 49783 | 56644 | 42730 |
| $R_{work}/R_{free}$ | 19.01/22.41 | 19.20/ 21.30 | 20.61/23.99 |
| Amino acids | 546 | 546 | 546 |
| Waters | 204 | 222 | 147 |
| Bonds (Å) | 0.008 | 0.009 | 0.008 |
| Angles (deg) | 0.94 | 0.92 | 0.94 |
| Average B factors compound(s) ($Å^2$) | 38.22 | 42.97 | 55.44 |
| Average B factors solvent ($Å^2$) | 43.17 | 46.81 | 60.12 |
| Ramachandran plot | | | |
| Favoured regions (%) | 97.38 | 97.42 | 97.2 |
| Allowed (%) | 2.06 | 2.03 | 2.43 |
| Outliers (%) | 0.56 | 0.55 | 0.37 |

| | Structure with fragment 4 | Structure with fragment 5 | Structure with fragment 6 and 10 |
|---|---|---|---|
| **Data collection** | | | |
| PDB id | 9QZU | 9QZV | 9QZW |
| Data collection date | April 2024 | April 2023 | April 2024 |
| Wavelength (Å) | 0.98 | 0.98 | 0.98 |
| Temperature (K) | 293 | 293 | 293 |
| a, b, c (Å) | 95.33, 95.33, 249.24 | 94.49, 94.49, 246.38 | 95.31, 95.31, 248.64 |
| α, β, γ (°) | 90, 90, 120 | 90, 90, 120 | 90, 90, 120 |
| Resolution range (Å) | 126.96-2.36 (2.36-2.40) | 123.19-2.04 (2.04-2.05) | 27.35-2.28 (2.28-2.32) |
| Total no. of reflections | 1226546 | 6788913 | 2278065 |
| No. of unique reflections | 52151 | 67981 | 57693 |
| Completeness (outer shell) (%) | 100 (100) | 100 (100) | 100 (100) |
| Redundancy (outer shell) | 23.52 (16.3) | 99.86 (67.3) | 39.49 (27.5) |
| I/σ(I) | 3.86(1.08) | 5.18 (0.89) | 4.15 (0.93) |
| $CC_{1/2}$ | 0.93 (0.44) | 0.96 (0.61) | 0.95 (0.50) |
| $R_{split}$ (%) | 24.59 (102.50) | 17.04 (67.15) | 19.52 (101.92) |
| Wilson B factor (Å$^2$) | 40.93 | 48.89 | 47.96 |
| Total no. of images | 34930 | 58401 | 22895 |
| Indexed images | 4325 | 10459 | 6330 |
| No. of supports collected | 2 | 4 | 2 |
| Soaking time (h) | 4:30-5h | 4:30-6h | 6h |
| Fragment(s) modelled | 1 | 1 | 2 |
| **Structure refinement** | | | |
| Resolution range (Å) | 2.36-27.35 | 2.04-29.1 | 2.28-27.35 |
| No. of reflections used in refinement | 28451 | 42833 | 31383 |
| $R_{work}/R_{free}$ | 20.61/25.74 | 20.08/22.97 | 20.12/23.96 |
| Amino acids | 546 | 546 | 546 |
| Waters | 147 | 142 | 136 |
| Bonds (Å) | 0.007 | 0.008 | 0.007 |
| Angles (deg) | 0.91 | 0.94 | 0.94 |
| Average B factors compound(s) (Å$^2$) | 50.8 | 57.88 | 56.92 |
| Average B factors solvent (Å$^2$) | 49.24 | 61.67 | 58.65 |
| Ramachandran plot | | | |
| Favoured regions (%) | 95.58 | 97.41 | 97.05 |
| Allowed (%) | 4.05 | 2.22 | 2.58 |
| Outliers (%) | 0.37 | 0.37 | 0.37 |

| | Structure with fragment 7 | Structure with fragment 8 | Structure with fragment 9 |
|---|---|---|---|
| **Data collection** | | | |
| PDB id | 9R02 | 9QZX | 9QZZ |
| Data collection date | Novemeber 2023 | April 2023 | Novemeber 2023 |
| Wavelength (Å) | 0.98 | 0.98 | 0.98 |
| Temperature (K) | 293 | 293 | 293 |
| a, b, c (Å) | 94.55, 94.55, 247.19 | 94.36, 94.36, 247.04 | 94.36, 94.36, 246.03 |
| α, β, γ (°) | 90, 90, 120 | 90, 90, 120 | 90, 90, 120 |
| Resolution range (Å) | 38.87-2.02 (2.02-2.06) | 123.52-1.95 (1.95-1.96) | 38.78-2.04 (2.04-2.08) |
| Total no. of reflections | 2985579 | 6366526 | 3191333 |
| No. of unique reflections | 81134 | 80801 | 78112 |
| Completeness (outer shell) (%) | 100 (100) | 100 (100) | 100 (100) |
| Redundancy (outer shell) | 36.80 (23.3) | 78.79 (52.6) | 40.86 (26.4) |
| I/σ(I) | 4.39 (0.96) | 4.74 (0.88) | 4.61 (0.86) |
| $CC_{1/2}$ | 0.95 (0.48) | 0.96 (0.62) | 0.95 (0.46) |
| $R_{split}$ (%) | 18.36 (102.40) | 17.23 (72.87) | 17.14 (108.73) |
| Wilson B factor (Å$^2$) | 35.02 | 44.84 | 36.78 |
| Total no. of images | 41093 | 25452 | 43066 |
| Indexed images | 6502 | 6337 | 8628 |
| No. of supports collected | 3 | 2 | 2 |
| Soaking time (h) | 5h | 5:30-6h | 4-4:30h |
| Fragment(s) modelled | 1 | 2 | 2 |
| **Structure refinement** | | | |
| Resolution range (Å) | 2.02-38.87 | 1.95-28.39 | 2.04-38.78 |
| No. of reflections used in refinement | 43784 | 49054 | 42187 |
| $R_{work}/R_{free}$ | 19.49/22.00 | 20.13/23.09 | 19.28/22.35 |
| Amino acids | 546 | 546 | 546 |
| Waters | 179 | 189 | 165 |
| Bonds (Å) | 0.008 | 0.008 | 0.008 |
| Angles (deg) | 0.94 | 0.92 | 0.95 |
| Average B factors compound(s) (Å$^2$) | 40.01 | 46.7 | 43.28 |
| Average B factors solvent (Å$^2$) | 46.17 | 51.36 | 50.07 |
| Ramachandran plot | | | |
| Favoured regions (%) | 97.56 | 96.68 | 97.24 |
| Allowed (%) | 2.25 | 2.95 | 2.39 |
| Outliers (%) | 0.19 | 0.37 | 0.37 |

**Table S3.** The resulting datasets binned according to resolution. The data display that the data-collection hit rate is reduced at lower resolution.

| Resolution range (Å) | No. of datasets | Average no. of indexable images | Average no. of total images | Average hit-rate | Average resolution (Å) |
|---|---|---|---|---|---|
| All | 96 | 7171 | 30253 | 24% | 2.23 |
| 1.81-2.0 | 24 | 9910 | 34965 | 29% | 1.96 |
| 2.01-2.2 | 44 | 7572 | 31160 | 26% | 2.10 |
| 2.21-2.5 | 14 | 5640 | 27996 | 22% | 2.35 |
| 2.51-4.3 | 14 | 2743 | 21585 | 13% | 3.01 |
